# ApoE ε4-dependent alteration of CXCR3^+^CD127^+^ CD4^+^ T cells is associated with elevated plasma neurofilament light chain in Alzheimer’s disease

**DOI:** 10.1101/2024.05.28.596276

**Authors:** Dan Hu, Mei Chen, Xuyang Li, Peter Morin, Sarah Daley, Yuyang Han, Martin Hemberg, Howard L. Weiner, Weiming Xia

**Author notes:** To whom correspondence should be addressed: Weiming Xia, Dan Hu or Howard L. Weiner.

## Abstract

Recent findings indicate a correlation between the peripheral adaptive immune system and neuroinflammation in Alzheimer’s disease (AD). To characterize the composition of adaptive immune cells in the peripheral blood of AD patients, we utilized single-cell mass cytometry (CyTOF) to profile peripheral blood mononuclear cells (PBMCs). Concurrently, we assessed the concentration of proteins associated with AD and neuroinflammation in the plasma of the same subjects. We found that the abundance of proinflammatory CXCR3^+^CD127^+^ Type 1 T helper (Th1) cells in AD patients was negatively correlated with the abundance of neurofilament light chain (NfL) protein. This correlation is apolipoprotein E (ApoE) ε4-dependent. Analyzing public single-cell RNA-sequencing (scRNA-seq) data, we found that, contrary to the scenario in the peripheral blood, the cell frequency of CXCR3^+^CD127^+^ Th1 cells in the cerebrospinal fluid (CSF) of AD patients was increased compared to healthy controls (HCs). Moreover, the proinflammatory capacity of CXCR3^+^CD127^+^ Th1 cells in the CSF of AD patients was further increased compared to HCs. These results reveal an association of a peripheral T-cell change with neuroinflammation in AD and suggest that dysregulation of peripheral adaptive immune responses, particularly involving CXCR3^+^CD127^+^ Th1 cells, may potentially be mediated by factors such as ApoE ε4 genotype.

**One sentence summary:** An apolipoprotein E (ApoE) ε4-dependent alteration of CD4 T cell subpopulation in peripheral blood is associated with neuroinflammation in patients with Alzheimer’s disease.

## INTRODUCTION

Alzheimer’s disease (AD) is a progressive and debilitating neurodegenerative condition affecting the central nervous system (CNS). Neuroinflammation in AD is characterized by innate immune responses involving CNS-resident reactive astrocytes and activated microglia surrounding amyloid plaques, implicating their role in disease pathogenesis (*1–4*). Investigation into the role of adaptive immunity and the peripheral immune system in AD is ongoing. Animal studies suggest a functional link between the peripheral immune system and the CNS in AD, implying that the peripheral immune system, including T cells, might influence AD progression (as reviewed by Cao, et al. in (*5*)). Recent studies have shown that in various mouse AD models employing distinct methodologies, brain T cells, stemming from microglia-mediated peripheral T cell migration to the brain, exhibit divergent effects, being identified as either detrimental (*6*) or protective (*7*) within their respective model contexts. In humans, alterations in T cell phenotypes in peripheral blood have been associated with AD. An increased CD45RO/CD45RA ratio in CD4^+^ T cells in the blood was found in AD patients or those with cognitive abnormalities compared to age-matched non-dementia controls (*8*), suggesting heightened CD4^+^ T cell activities in these individuals. Additionally, increased proportions of activated HLA-DR^+^ CD4^+^ and CD8^+^ T cells were observed in the blood of patients with mild cognitive impairment (MCI) or mild AD, correlating with the neuropsychological deficits typically found in these patients (*9*). Increased CD45RA^+^CD27^−^ effector memory CD8^+^ T (CD8 T_EMRA_) cells were detected in both blood and CSF in AD, showing a negative association with cognitive function (*10*). Mitogenic activation of CD4^+^ and CD8^+^ T cells derived from the blood of patients with sporadic late-onset AD displayed increased tyrosine phosphorylation, CD69 expression, and proliferation (*11*). Moreover, reduced PLCγ2 activity was reported in peripheral blood mononuclear cells (PBMCs), including CD4^+^ and CD8^+^ T cells, in AD (*12*), which is a downstream target of receptor tyrosine kinases. Additionally, AD patients were reported to harbor amyloid-β42-specific T-cells in the blood (*13, 14*).

The immune cell profiling studies conducted on peripheral blood in patients with AD have typically concentrated on a limited array of cell types characterized by predetermined cell surface markers (*15*). However, a recent study capitalized on high-parameter mass cytometry by time-of-flight (CyTOF) to investigate the responses of peripheral immune cells from individuals with AD or Parkinson’s disease (PD) to seven established immune stimulants. This study utilized an antibody panel recognizing 12 cell surface markers and 15 intracellular signaling molecules (*12*). By categorizing immune cell subtypes based on classic, pre-defined cell surface markers, this study unveiled altered PLCγ2, Stat1 and Stat5 signaling in AD, albeit without significant changes in the abundance of immune cell subtypes between AD patients and HCs. The divergence between these findings and the previous reports (*8–10*) necessitates further exploration to reconcile the observed differences.

In the present study, we used CyTOF with an antibody panel recognizing 29 immune cell surface markers to profile peripheral blood mononuclear cells (PBMCs) from individuals with AD. We utilized a dimensionality reduction and clustering approach to analyze the data, aiming to identify non-pre-defined cell populations exhibiting altered abundance in AD patients. Additionally, we conducted assessments of blood AD-associated proteins in the plasma obtained from the same individuals using enzyme-linked immunosorbent assay (ELISA). Furthermore, we analyzed publicly available single-cell RNA-sequencing (scRNA-seq) datasets obtained from both healthy individuals and AD patients, focusing on the cell populations that exhibited alterations identified in AD patients through CyTOF analysis. Our investigation unveiled alterations in CD4^+^ T cell subpopulations within both the peripheral blood and CSF of individuals with AD, in particular, the frequency of CD161^+^CCR6^+^ Th17 subset was increased in peripheral blood of AD patients and an apolipoprotein E (ApoE) ε4-dependent reversed correlation between peripheral CXCR3^+^CD127^+^ Th1 cells and plasma neurofilament light chain (NfL) in AD patients was revealed.

## RESULTS

### Overview of the experimental and analytical design

The study aimed to identify correlations between AD and specific peripheral blood cell populations. **Figure 1A** illustrates an overview of the study’s structure. In summary, we performed immunophenotyping of PBMCs obtained from 18 AD patients and 19 non-dementia (ND) controls (**Supplementary Tables 1 and 2**) using CyTOF to assess the expression of 29 immune cell surface markers (**Supplementary Table 3**). The data files underwent two types of analyses: manual gating and dimensionality reduction with clustering. Manual gating involved analyzing individual data files for all 37 samples separately, following a defined sequential gating strategy based on pre-established cell surface markers. On the other hand, for dimensionality reduction and clustering analyses, we selected 32 samples (AD: n = 16; ND: n = 16) with the highest event numbers. We randomly sampled an equal number of cells (38,000 CD45^+^ cells or 12,000 CD3^+^ cells per sample) from each of these samples and merged them into a single file, termed the ‘Merged-PBMC File’ or ‘Merged T-cell File.’ This approach ensured an equitable representation of each individual’s cells in the merged file, facilitating equal representation and statistical analysis.

**Figure 1.**
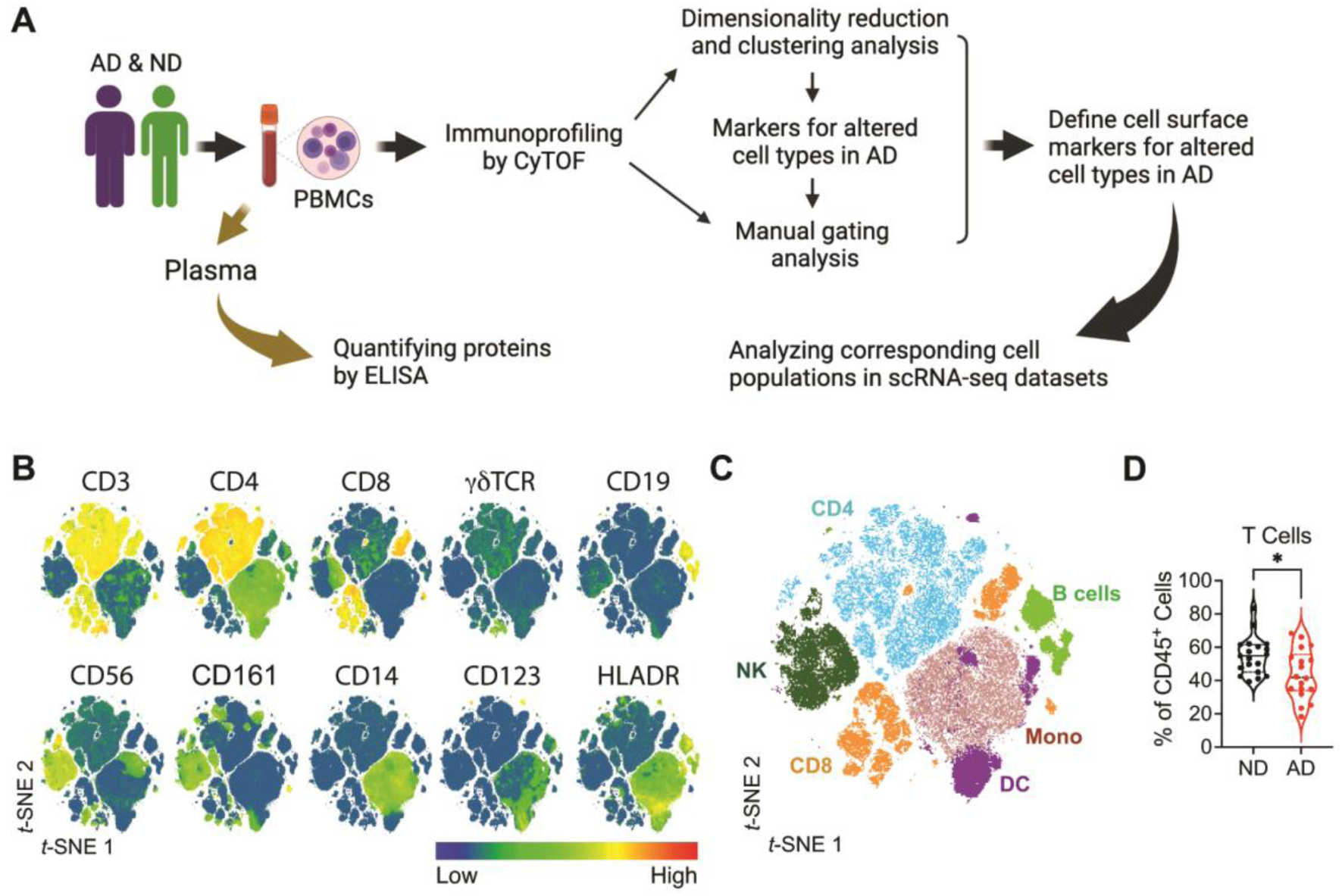
Reduced total T cell abundance in the peripheral blood of AD patients. **A)** Scheme for overall experimental and analytical workflow. Peripheral blood was drawn from patients with AD and non-dementia (ND) controls, plasma was collected for protein quantification by ELISA, and PBMCs were isolated for immune cell profiling by mass cytometry (CyTOF). Cell surface markers defined by CyTOF analysis for the altered cell populations in AD were used to identify the corresponding cell populations in a scRNA-seq dataset. The gene expression profiles of these cell populations were extracted and subjected to computational analysis for relevant gene expression features and cell signaling pathways. **B-D**) Reduced total T cell abundance in the peripheral blood of AD patients. PBMCs were isolated from non-dementia controls (ND) and patients with AD (AD) for CyTOF analysis. **B**) Expression of immune cell surface markers. tSNE plot was generated using the Merged PBMC File (38,000 CD45^+^ cells per individual, and a total of 1,216,000 cells) (ND, n = 16; AD, n = 16) using the expression information of the 29 cell surface markers. **C**) A distribution map of main immune cell populations. **D**) Total T cell frequency comparisons between controls and patients. \**p* < 0.05; ns, not significant (Mann-Whitney test; mean ± s.d.).

We next utilized manual gating analysis to delineate the cell surface markers of altered cell clusters identified in AD through dimensionality reduction, and analyzed the association between cell frequencies and AD/ND status. Additionally, plasma samples from the same individuals were collected and analyzed for the protein concentration of IL-2, IFN-γ, Tau, pTau181, and NfL using ELISA. We further analyzed the relation between the cell population frequencies and plasma protein concentrations within AD and ND cohorts respectively and contrasted the partial correlation between the two groups. Finally, we investigated the cell populations of interest using publicly available single-cell RNA-sequencing (scRNA-seq) datasets on PBMCs (GSE158055) (*16*) and CSF cells (GSE134578) (*10*). Leveraging the cell surface markers defined in the CyTOF analysis for the altered AD clusters, we identified corresponding cell clusters in healthy subjects within the scRNA-seq datasets. Subsequently, we extracted the gene expression profiles of these identified cell clusters to define their gene expression features and infer their functional characteristics through gene set enrichment and pathway analyses.

### Reduced total T cell abundance in the peripheral blood of patients with AD

To delineate six primary immune cell populations in peripheral blood, we initially employed a sequential gating strategy (manual gating). These populations encompassed CD4^+^ T cells (CD45^+^CD3^+^TCRγδ^−^CD4^+^CD8^−^), CD8^+^ T cells (CD45^+^CD3^+^TCRγδ^−^CD4^−^CD8^+^), B cells (CD45^+^CD14^−^CD66b^−^CD3^−^ CD19^+^), NK cells (CD45^+^CD14^−^CD66b^−^CD3^−^CD19^−^CD20^−^CD123^−^CD56^+^CD161^+^), dendritic cells (CD45^+^CD14^−^CD66b^−^CD3^−^CD19^−^CD20^−^CD56^−^HLADR^+^), and monocytes (CD45^+^CD66b^−^CD3^−^CD19^−^CD14^+^). We visualized both the expression of lineage markers (**Fig. 1B**) and the six cell populations (**Fig. 1C**) within the t-Distributed Stochastic Neighbor Embedding (tSNE) plot derived from the Merged-PBMC File. Through manual gating analysis of individual CyTOF data files, we evaluated the frequency of these primary immune cell populations within the PBMCs of AD patients and ND donors. While no significant differences emerged in CD4^+^ T cells, CD8^+^ T cells, B cells, NK cells, dendritic cells, and monocytes (**Supplementary Fig. 1**), a reduction in the frequency of total T cells (CD45^+^CD3^+^) was evident in AD patients (**Fig. 1D**).

### Dimensional analysis of immune cell components in the peripheral blood in AD

To analyze highly multiplexed datasets and unveil novel or unexpected cell subtypes while preserving valuable information, we employed a data-driven, open-ended dimensionality reduction and clustering approach to analyze CyTOF data (*17, 18*). Utilizing the FlowSOM analysis platform (*19*), we processed the Merged-PBMC File, employing a self-organizing map (SOM) to identify thirty-four clusters visualized in the tSNE plot of the Merged-PBMC File (**Fig. 2A**). We computed the percentage of events in each cluster for all individuals and assessed their statistical significance between AD and ND samples. Notably, the frequencies of cells in Clusters 19 and 27 exhibited significant reduction in AD patients (**Fig. 2B & C**). Cluster 27 cells predominantly clustered within the CD4^+^ T cell area, whereas Cluster 19 cells were more diffusely distributed around the T cell region in the tSNE plot (**Fig. 2D**). The heat map generated based on individual cluster marker expression revealed distinctive characteristics: Cluster 19 cells were CD3^+^CD4^−^CD8^−^CD45RA^−^CD45RO^low^ while Cluster 27 cells were CD3^+^CD4^+^CD45RA^−^CD45RO^+^CXCR3^+^CD127^+^CCR7^+^CD161^−^CCR6^−^ (**Fig. 2E**). CD45RA^−^, CD45RO^+^, CCR7^+^ and CXCR3^+^ are characteristic features of central memory Th1 cells (*20*). Consequently, we identified a diminished abundance of CD4 and CD8-double negative (DN) T cells, alongside reduced CXCR3^+^CD127^+^ central memory Th1 cells in the peripheral blood of AD patients.

**Figure 2.**
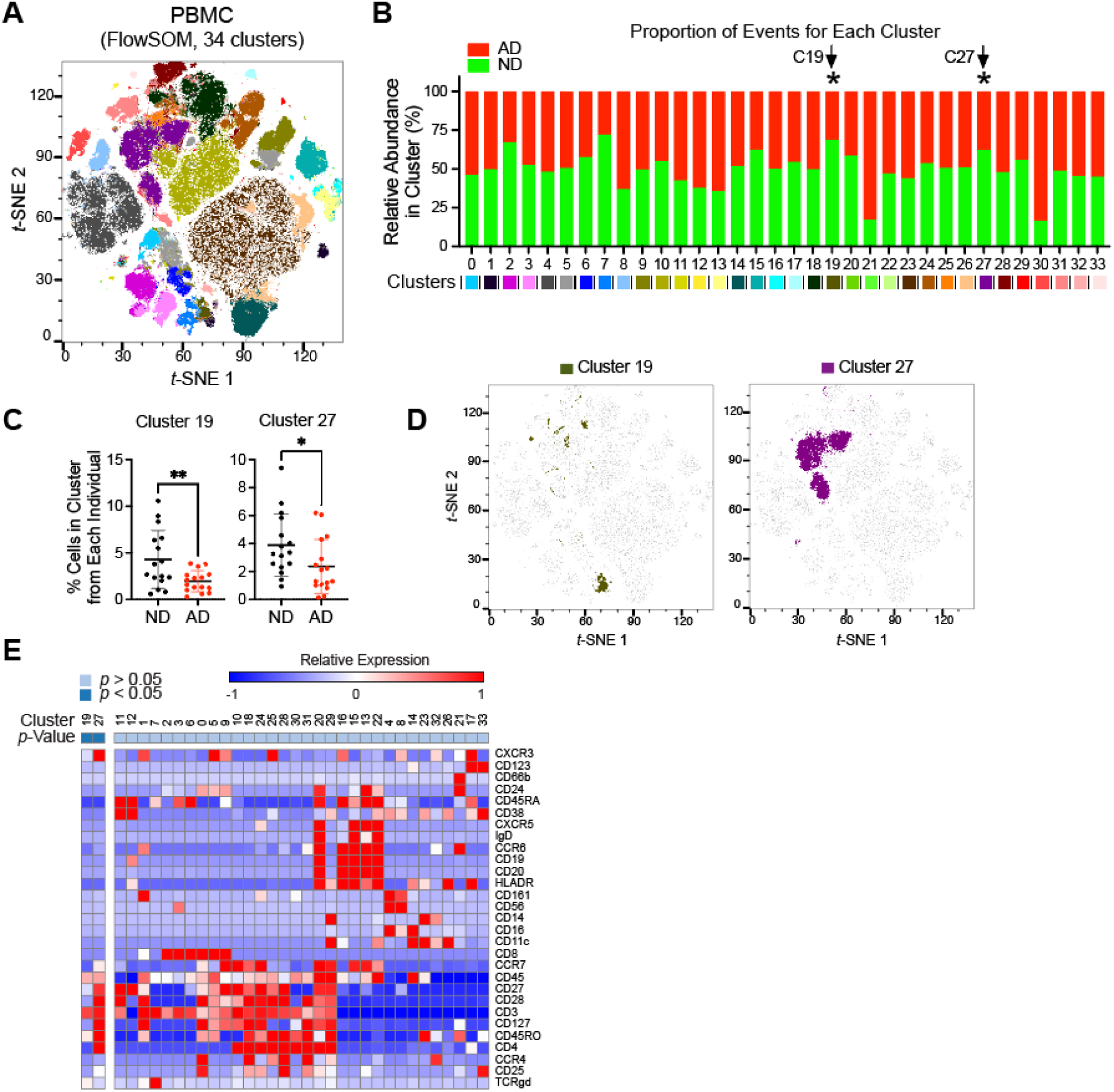
Dimensional analysis of PBMCs identifies reduced T cell subsets in AD. FlowSOM analysis with the tSNE-plot of the Merged PBMC File using the expression information of the 29 cell surface markers. **A**) Thirty-four clusters (color coded) were identified in the tSNE plot. **B**) Stacked bars show the respective abundance of cells from NDs and AD patients for each cluster. The color codes for the clusters match with that in the tSNE plot. The percentage of cells contributed by each individual in each cluster was used to calculate the statistical difference between controls and patients, and significant difference was found in Clusters 19 and 27 (C19 and C27) (\**p* < 0.05, two-tailed, unpaired *t*-test). **C**) The percentage of cells contributed by each individual in Clusters 19 and 27 (indicated by dots) was significantly reduced in AD patients compared to controls (mean ± s.d.). **D**) Distribution of Clusters 19 and 27. **E**) Heatmap depicting per cluster median staining intensity of the 29 cell surface markers on all 32 samples from CyTOF analysis.

### Dimensional analysis of T cell subpopulations in the peripheral blood in AD

To delve deeper into the association between T cell subsets and AD, we randomly sampled an equal number of T cells (12,000 CD3^+^ cells per sample) from the 32 samples (AD, n = 16; ND, n = 16), amalgamating them into a single file containing 384,000 cells, termed the ‘Merged-T-cell File.’ Employing the FlowSOM analysis platform, we conducted a 34-cluster analysis on the Merged-T-cell File, visualizing them using a tSNE plot (**Fig. 3A**). We then computed the percentage of events within each cluster for all individuals, and we assessed the difference between AD and ND samples. Notably, the frequencies of cells in Clusters 8, 20, and 27 exhibited significant increases in AD patients, whereas the frequency of Cluster 12 cells was significantly reduced (**Fig. 3B & C**). Visualizing CD4^+^, CD8^+^, CD4^+^CD8^+^ and CD4^−^CD8^−^ T cells within the tSNE plot (**Fig. 3D**), we observed that all four altered clusters were localized within the CD4^+^ T cell region (**Fig. 3E**). The heat map, based on individual cluster marker expressions, revealed distinctive characteristics: all four altered clusters were CD4^+^CD8^−^CD45RA^−^CD45RO^+/low^, with Cluster 12 showing a remarkably similar cell surface marker expression pattern to that of Cluster 27 cells in **Fig. 2E**. Cluster 8 exhibited a CD27^+^CCR7^+^CD28^+^ profile reminiscent of central memory T cells, while Cluster 20 displayed a CCR4^+^CD27^−^CCR6^−^ profile akin to Th2 cells, and Cluster 27 exhibited a CCR6^+^CD161^+^ profile reminiscent of Th17 cells (**Fig. 3F**). Thus, through high-dimensional analysis of total T cells we identified four altered effector/memory CD4^+^ T cell subpopulations in the peripheral blood of AD patients.

**Figure 3.**
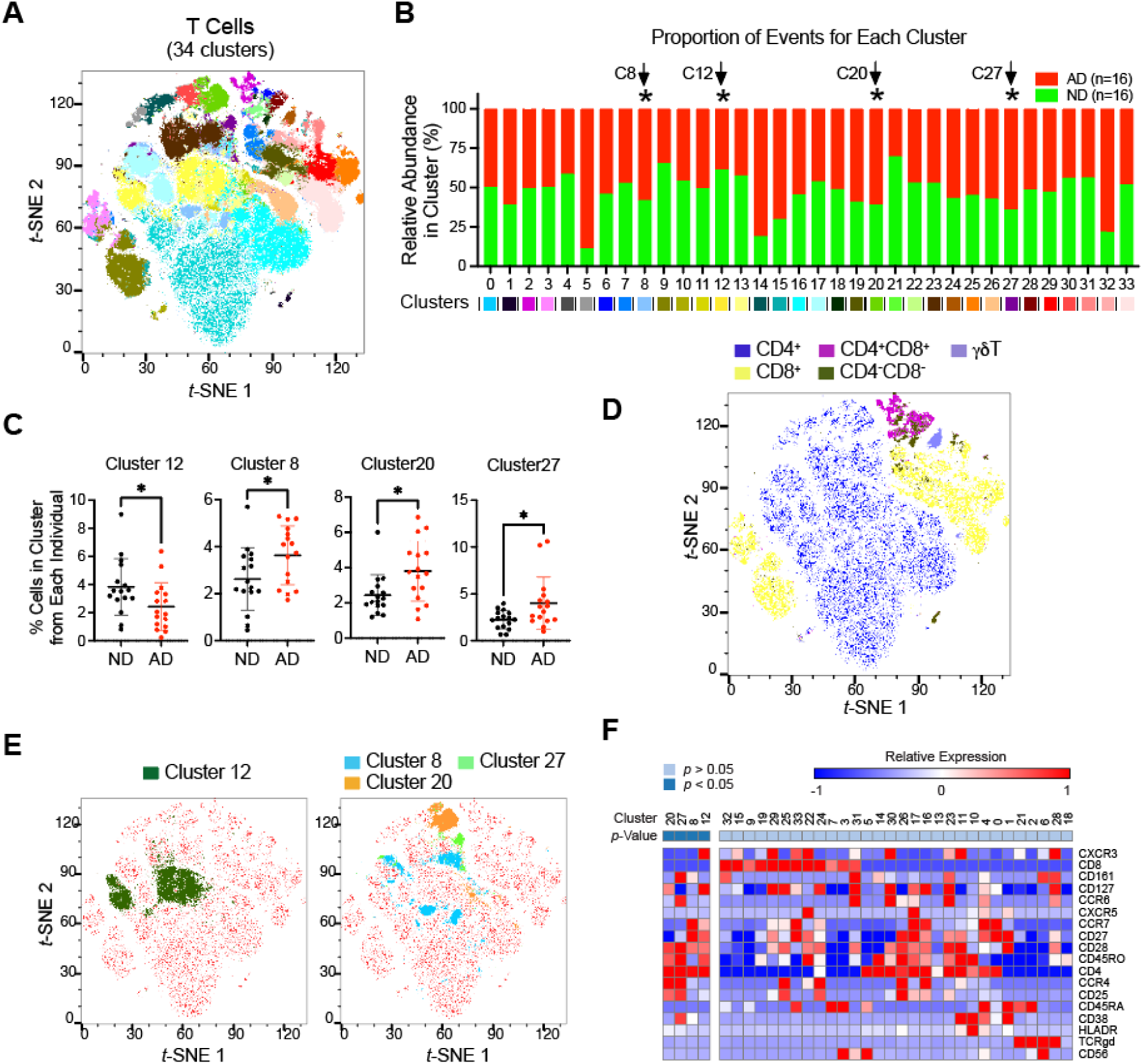
Dimensional analysis of peripheral T cells identifies altered T cell subpopulations in AD. FlowSOM analysis with the tSNE-plot of the Merged T-cell File using the expression information of the 29 cell surface markers. **A**) Thirty-four clusters (color coded) were identified in the tSNE plot. **B**) Stacked bars show the respective abundance of cells from NDs and AD patients for each cluster. The color codes for the clusters match with that in the tSNE plot. The percentage of cells contributed by each individual in each cluster was used to calculate the statistical difference between controls and patients, and significant difference was found in Clusters 8, 12, 20 and 27 (C8, C12, C20 and C27) (\**p* < 0.05, two-tailed, unpaired *t*-test). **C**) The abundance of Cluster 12 was reduced in AD while increased for Clusters 8, 20 & 27 (individuals indicated by dots, mean ± s.d.). **D**) A distribution map of main T cell populations. **E**) Clusters 12, 8, 20 and 27 were located within the CD4^+^ T cell population. **F**) Heatmap depicting per cluster median staining intensity of T cell relevant cell surface markers on all 32 samples from CyTOF analysis.

### Increased peripheral Th17 subpopulation in patients with AD

The FlowSOM-generated clusters were based on 29 cell surface markers, which proved excessive for practical cell subset categorization. Moreover, the cell surface marker expressions depicted in heat maps (**Fig. 2E** and **Fig. 3F**) median expression levels across cells within individual clusters, lacking resolution on specific marker positivity or negativity within a cluster at the cell level. We next utilized manual gating analysis to delineate the cell surface markers of these altered T-cell clusters among the 18 AD patients and 19 ND controls, aiming to validate the outcomes derived from our clustering analyses. Hence, employing a sequential gating strategy, we identified a reduced set of consistently expressed or absent markers to define the double-negative T cells and four subpopulations of effector/memory CD4^+^ T cells identified in dimensional and clustering analysis: CD4^−^CD8^−^ (Cluster 19 of PBMCs, DN), CXCR3^+^CD127^+^ (Cluster 27 of PBMCs and Cluster 12 of T cells, Th1-subset), CCR4^+^CD27^−^ (Cluster 20 of T cells, Th2-subset), CCR7^+^CD27^+^CD45OR^low^CD161^+^ (Cluster 8 of T cells, central memory (cm)-subset) and CD161^+^CCR6^+^CCR4^+^CD127^−^ (Cluster 27 of T cells, Th17-subset) T cell subpopulations (**Fig. 4A**). Given the age difference between AD and ND groups and uneven distribution of male and female in the cohorts, we adopted the multivariate regression model adjusted by age and sex regarding AD and ND controls to assess the cell frequency change. Results showed that among the PBMCs, the cell frequency change between AD and ND controls for DN T cells was insignificant (**Supplementary Table 4**). However, the average cell frequency of CD161^+^CCR6^+^CCR4^+^CD127^−^ Th17-subset was increased from the 0.5% in control group to 0.71% in AD patients with *p*-value of 0.0327 and FDR adjusted *p*-value of 0.131 (**Fig. 4B & C**). Our results also suggest the trend of increased CCR4^+^CD27^−^ Th2-subset cells and decreased CXCR3^+^CD127^+^ Th1-subset cells in the peripheral blood of AD patients (**Fig. 4B** and **Supplementary Table 4**).

**Figure 4.**
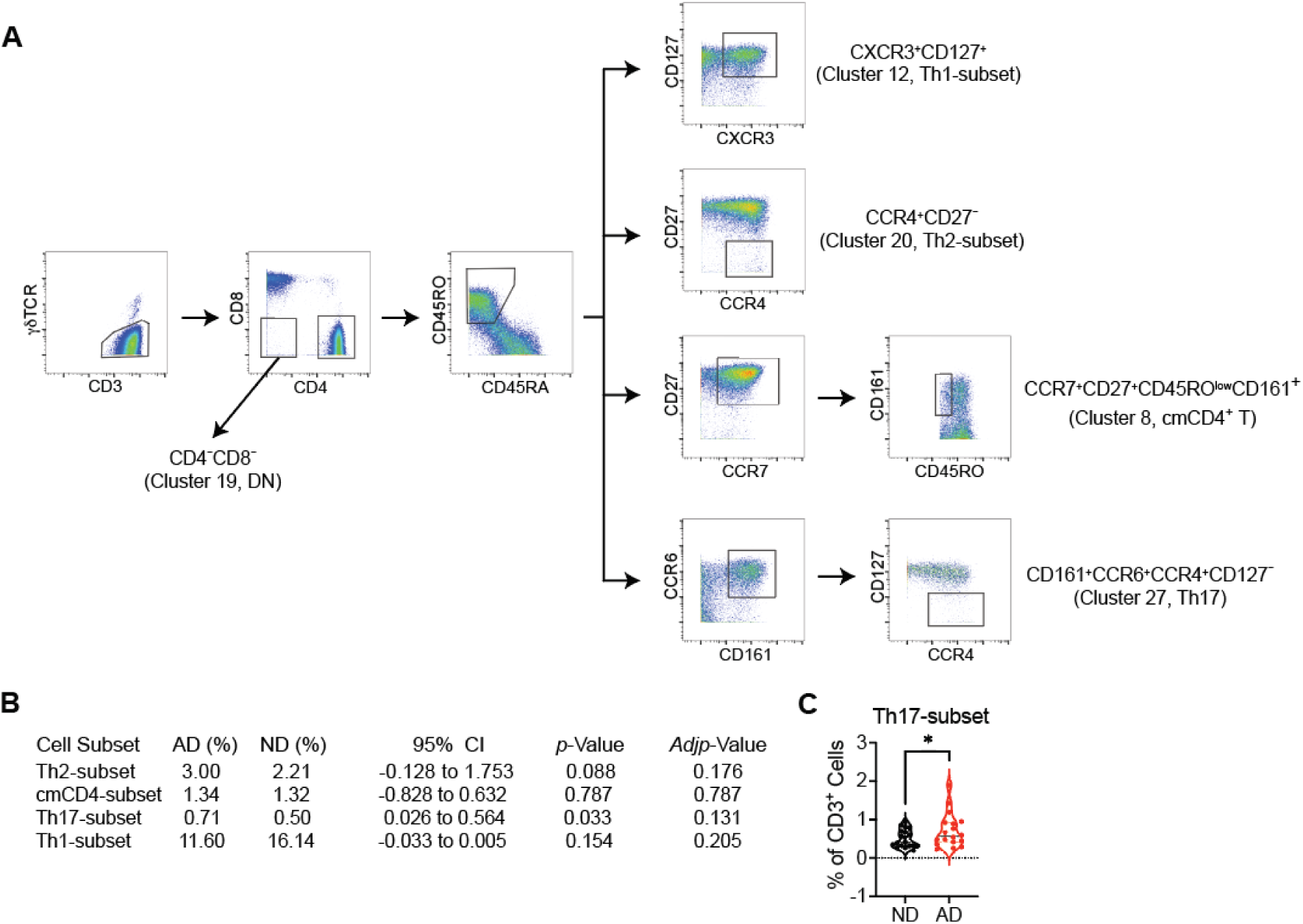
Increased CD161^+^CCR6^+^CCR4^+^CD127^−^ (Th17-subset) T cells in patients with AD. **A**) Manual gating strategy for the altered T cell subpopulations identified with dimensional analysis. **B**) Multivariate regression model of the cell frequencies (%) of CD4^+^ T cell subsets within the total T cell population, adjusted for age and sex. To correct for multiple testing, FDR adjusted *p*-values are shown. **C**) Manually gated CD161^+^CCR6^+^CCR4^+^CD127^−^ (Th17-subset) T cells were significantly increased in patients with AD. \**p* < 0.05 (multivariate regression adjusted by age and sex). Each dot represents an individual (ND, n = 19; AD, n =18).

### ApoE ε4-dependent inverse correlation between peripheral CXCR3^+^CD127^+^ Th1 cells and plasma NfL levels in AD patients

The microtubule-associated protein tau (encoded by *MAPT*) is abundant in CNS neurons (*21, 22*). It becomes hyperphosphorylated and forms insoluble neurofibrillary tangles in brains of patients with AD and other neurodegenerative diseases (*23–25*). Plasma tau is elevated in AD patients (*26, 27*). Plasma tau phosphorylated at residue 181 (pTau181) is correlated with tau pathology associated with AD (*28, 29*). Neurofilament light chain (NfL) levels are increased in the blood of AD patients and are associated with AD diagnosis and cognitive decline (*30–33*). Our ELISA analysis confirmed significantly higher plasma levels of Tau, pTau181, and NfL proteins in AD patients compared to ND donors (**Fig. 5A**). Partial within-group correlation analysis controlling for age and sex did not identify any significant correlations between these T cell subsets and plasma NfL levels, in neither the AD group nor in ND control group (**Supplementary Table 5**) when controlled for age and sex. However, when the analysis was controlled for the presence of one or two copies of the ε4 variant of the human *APOE* (*APOE4*) gene, a known risk factor for AD that was associated with AD status in our cohort, in addition to age and sex, a reversed correlation between the cell frequency of CXCR3^+^CD127^+^ Th1-subset and plasma NfL levels was revealed in the AD group but not the ND control group (**Table 1**, **Supplementary Table 6** and **Fig. 5B**). We did not detect reduced plasma IFN-γ levels in AD patients (**Supplementary Fig. 2A**). CXCR3 is a chemokine receptor preferentially expressed on IFN-γ-expressing Th1 cells (*34*). CD127 is a receptor for IL-7 and blocks apoptosis during T cell differentiation and activation (*35–37*). Therefore, in this study, we defined CXCR3^+^CD127^+^ Th1 cells as a category of CD4^+^ T cells. Taken together, our results suggest an apolipoprotein E (ApoE) ε4-dependent reversed correlation between peripheral CXCR3^+^CD127^+^ Th1 cells and plasma NfL in AD patients.

**Figure 5.**
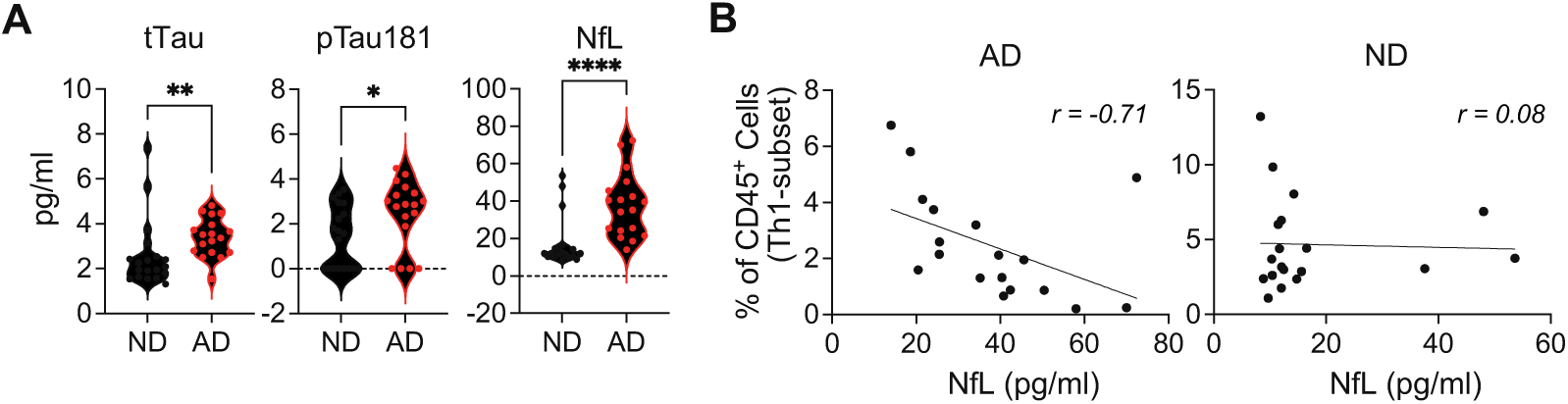
Peripheral abundance of CXCR3^+^CD127^+^ Th1 cells invertedly correlates with plasma NfL in AD patients. **A**) Quantification of plasma Tau, pTau181 and NfL protein levels by ELISA. \**p* < 0.05, \*\*\*\**p* < 0.0001 (Mann-Whitney test, mean ± s.d.). **B**) Inverted correlation between peripheral CXCR3^+^CD127^+^ Th1 cells and plasma NfL level in AD patients but not in ND controls (Partial Pearson’s correlation). Each dot represents an individual (ND, n = 19; AD, n =18).

**Table 1.** Partial Pearson’s correlation coefficient of cell population frequency versus NfL levels controlling for age, sex and *APOE4*.

| Cell subset | Correlation |  | 95% CI | AD |  |
| --- | --- | --- | --- | --- | --- |
|  | AD | ND |  | *p*-Value | *Adj**p*-Value |
| Th2-subset | 0.45 | -0.18 | (-0.083, 1.15) | 0.081 | 0.108 |
| cmCD4-subset | -0.0069 | 0.21 | (-0.85, 0.49) | 0.57 | 0.57 |
| Th17-subset | -0.69 | -0.12 | (-1.09, 0.02) | 0.056 | 0.108 |
| Th1-subset | -0.67 | 0.065 | (-1.22, -0.098) | 0.022 | 0.088 |

### CXCR3^+^CD127^+^ Th1 cells exhibit proinflammatory and cytotoxic gene signatures

To examine the global gene expression of CXCR3^+^CD127^+^ Th1 cells, we utilized a public scRNA-seq dataset (GSE158055) (*16*), extracting gene expression profiles from PBMCs of 5 healthy donors and integrating these samples to produce a total of 46,683 single-cell transcriptomes. The gene expression profiles of CD4^+^ T cells were obtained by initially selecting *CD3E*^+^*CD8A*^−^*CD4*^+^*TRDC*^−^ clusters from PBMCs, followed by a two-step algorithmic clustering to filter out CD8A^+^ clusters (CD8^+^ T cells) (**Methods and Supplementary Fig. 3**). Subsequently, the resulting CD4^+^ T-enriched population was clustered into 13 groups. Among these, we identified one cluster (Cluster 11) of *CD3E*^+^*CD4*^+^*CXCR3*^+^*IL7R*(CD127)^+^ cells (**Fig. 6A**) as CXCR3^+^CD127^+^ Th1 cells, constituting approximately 6% of the total CD3E^+^CD4^+^ cells (**Fig. 6B**). Compared to other CD4^+^ T cells, CXCR3^+^CD127^+^ Th1 cells exhibited elevated expression levels of lytic proteins (*GZMA* and *GZMK*), a proinflammatory chemokine (*CCL5*), a cell trafficking-mediating chemokine receptor (*CXCR3*), and a cell cytotoxicity regulator (**Fig. 6C**). Additionally, they displayed a significant enrichment in the molecular signature associated with CD4^+^ cytotoxic T cells (**Fig. 6D**). Ingenuity Pathway Analysis (IPA) revealed activation not only of multiple proinflammatory signaling pathways (cytokine storm, multiple sclerosis, and neuroinflammation signaling) in CXCR3^+^CD127^+^ Th1 cells but also enhancement in signaling pathways related to cell proliferation, activation, and migration (CDC42, PKCθ, and integrin signaling) (**Fig. 6E**). Thus, the global gene expression analysis showed that CXCR3^+^CD127^+^ Th1 cells were highly proliferative, cytolytic, proinflammatory, and prone to tissue homing.

**Figure 6.**
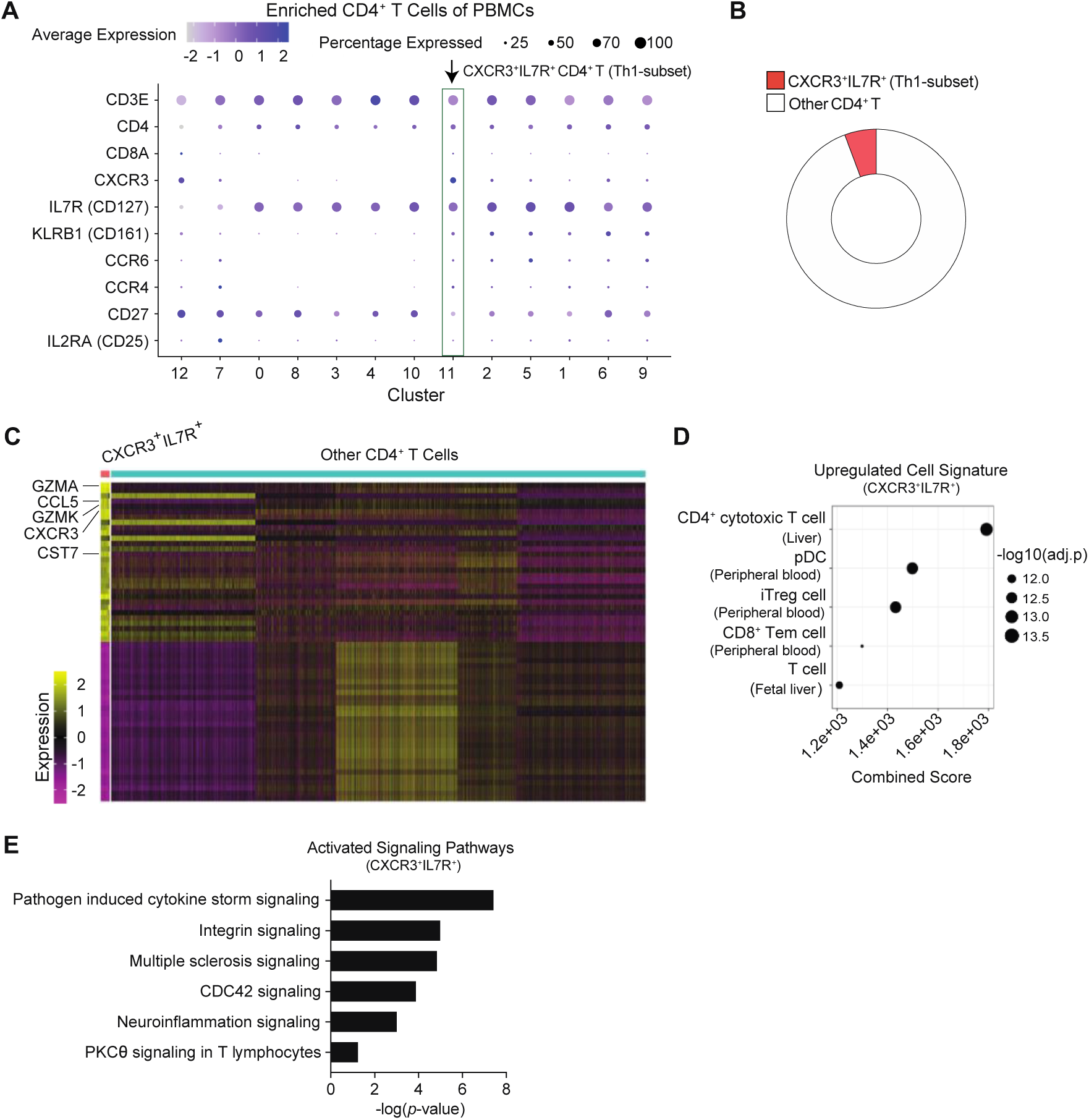
CXCR3^+^CD127^+^ Th1 cells are highly proinflammatory and cytotoxic. The gene expression features of CXCR3^+^CD127(IL7R)^+^ Th1 cells were analyzed as described in Materials and Methods using expression profiles of a CD4^+^ T cell-enriched population extracted from a public single-cell RNA-seq (scRNA-seq) dataset on PBMCs (GSE158055). **A**) Marker genes for CXCR3^+^CD127^+^ Th1 cells are shown. Green rectangle indicates a cluster enriched for CXCR3^+^CD127^+^ Th1 cells (Cluster 11). **B**) Frequency of CXCR3^+^CD127^+^ Th1 cells in total T cells was calculated with the number of single-cell transcriptomes in each cluster. **C**) Heatmap for the top 30 upregulated and top 30 down-regulated genes in CXCR3^+^CD127^+^ Th1 cells (adjusted *p* < 0.05, absolute fold-change (FC) ≥ 1.5). Five representative upregulated genes are shown ranked from the top by fold-change from high to low. **D**) Enrichr CellMarker Augmented 2021 analysis on the upregulated genes. Top five matched cell types with the lowest adjusted *p*-values are shown. **E**) IPA Canonical Pathway Analysis on all differentially expressed genes. Among eighteen signaling pathways with absolute Z-Score ≥ 2, six T cell-relevant pathways were selected and ranked from the top by -log(*p*-value) high to low.

### Elevated cell abundance and inflammatory potential of CXCR3^+^CD127^+^ Th1 cells in the CSF of AD patients

To investigate CXCR3^+^CD127^+^ Th1 cells in the CSF of AD patients, we utilized a public scRNA-seq dataset consisting of cells isolated from the CSF of 4 AD patients and 9 healthy controls (HC) (GSE134579) (*10*). Integration of all samples yielded 17,886 single-cell transcriptomes, comprising 5,073 from AD patients and 12,813 from HCs. Gene expression profiles of CD4^+^ T cells were obtained using a similar approach as described above (**Methods and Supplementary Fig. 4**). The resulting CD4^+^ T-enriched population was clustered into 8 clusters. Clusters 0, 1, 2, 3, and 4 were identified as *CD3E*^+^*CD4*^+^*CXCR3*^+^*IL7R*(CD127)^+^ cells (**Fig. 7A**), denoted as CXCR3^+^CD127^+^ Th1-enriched cells. In contrast to peripheral blood, where CXCR3^+^CD127^+^ Th1 cells constituted a minor subset of CD4^+^ T cells, these cells dominated the CD4^+^ T population in the CSF of HCs, comprising ∼84% of CD4^+^ T cells (**Fig. 7B, left**). In AD patients, the frequency of CXCR3^+^CD127^+^ Th1 cells was significantly increased, accounting for ∼96% of CD4^+^ T cells (**Fig. 7B, right**) (χ^2^ = 189.22, *p-value* < 2.2e-16, *φ* = 0.15). Differential gene expression analysis revealed 72 upregulated genes after multiple-testing correction, including proinflammatory genes *LTB* and *TNF*, and 9 downregulated genes in the CXCR3^+^CD127^+^ Th1 cells of AD patients compared to that of healthy controls. Pathway analysis revealed a significant increase in signaling pathways related to proinflammatory cytokine production (pathogen-induced cytokine storm signaling), cell death (death receptor signaling), and cell proliferation and activation (ILK, actin cytoskeleton, and Rho family GTPases signaling) in AD patients. Sirtuins are <u>n</u>icotinamide <u>a</u>denine <u>d</u>inucleotide (NAD)^+^-dependent histone deacetylases. The sirtuin signaling is important for anti-inflammatory and anti-apoptotic responses (*38*). Pathway analysis also revealed inhibited sirtuin signaling (**Fig. 7C**). Activation-induced cell death (AICD) is a critical self-restraining mechanism during T cell activation (*39*). Consequentially, activation of death receptor signaling and inhibition of anti-apoptotic responses in CXCR3^+^CD127^+^ Th1 cells of AD patients also suggested enhanced T cell activation. Collectively, these findings demonstrated the predominance of CXCR3^+^CD127^+^ Th1 cells exhibiting heightened proinflammatory capabilities in the CSF of AD patients.

**Figure 7.**
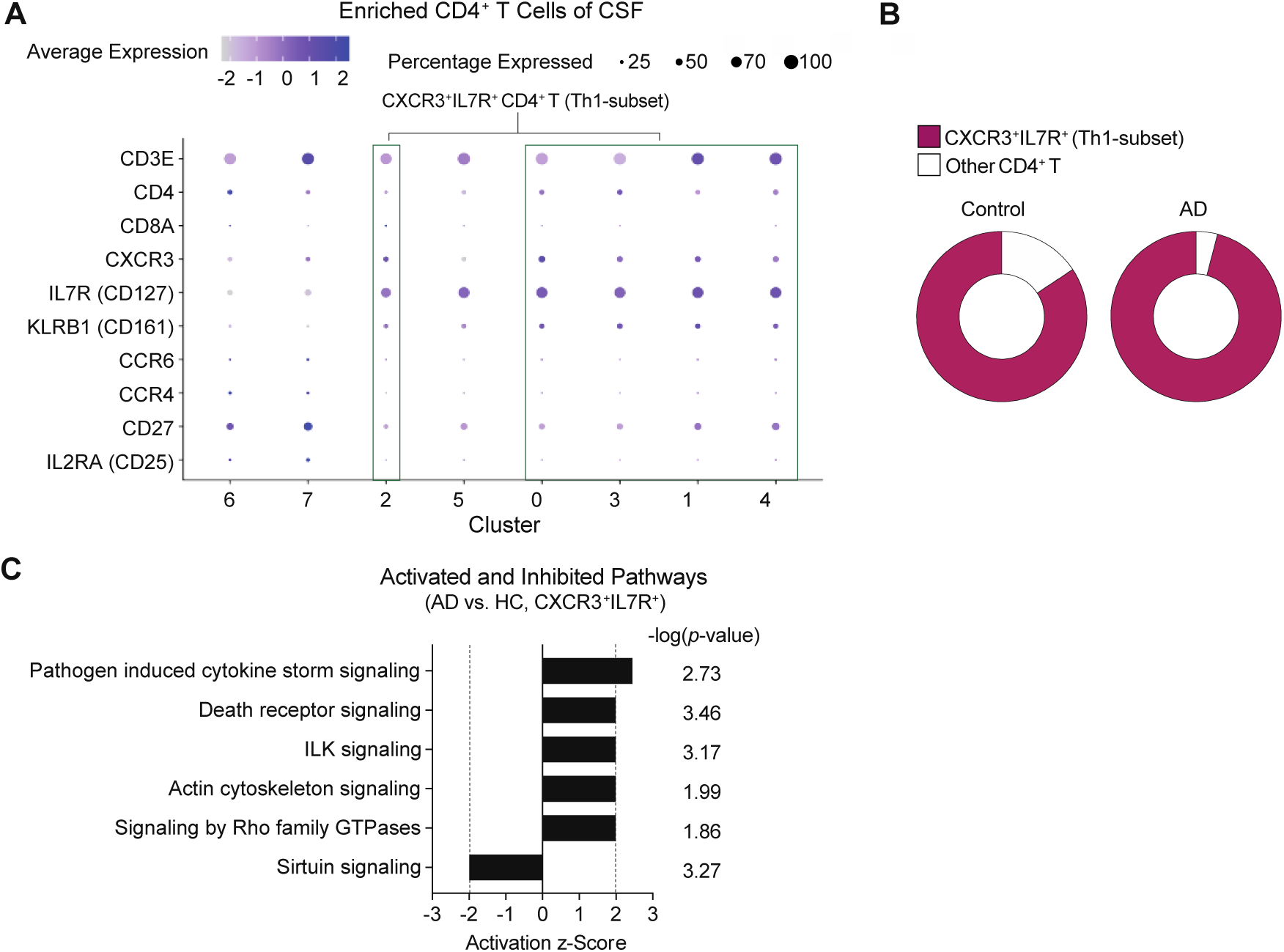
Increased CXCR3^+^CD127^+^ Th1 cells in the CSF of AD patients with elevated proinflammatory capacity. CXCR3^+^CD127(IL7R)^+^ Th1 cells in the CSF of AD patients were analyzed as described in Materials and Methods using expression profiles of a CD4^+^ T cell-enriched population extracted from a public single-cell RNA-seq (scRNA-seq) dataset (GSE134579). **A**) Marker genes for CXCR3^+^CD127^+^ Th1 cells are shown. Green rectangles indicate clusters enriched for CXCR3^+^CD127^+^ Th1 cells. **B**) Frequency of CXCR3^+^CD127^+^ Th1 cells in total T cells was calculated with the number of single-cell transcriptomes in each cluster. Difference in frequency among AD and HC CD4^+^ T cells was assessed with Chi-square test (χ^2^ = 189.22, *p-value* < 2.2e-16, *φ* = 0.15). **C**) IPA Canonical Pathway Analysis on all differentially expressed genes (adjusted *p* < 0.05). Among fourteen signaling pathways with absolute Z-Score ≥ 2, six T cell-relevant pathways were selected and ranked from the top by Z-score from high to low.

## DISCUSSION

Manual gating analysis with pre-defined cell surface markers for established cell types is a classical approach to analyze flow cytometric data as well as CyTOF data, which is a focused and targeted analysis based on prior biological knowledge. However, with dozens of simultaneously acquired parameters of CyTOF datasets available, dimensionality reduction and clustering analyses not only resolve the huge challenge of manually analyzing these highly multiplexed datasets, but also presents an open-ended strategy to identify unanticipated cell populations with biological significance (*17, 18*). Integrating these two approaches, we identified increased abundance of CD45RO^+^CD161^+^CCR6^+^CCR4^+^CD127^−^ CD4^+^ T cells in the peripheral blood of patients with AD as well as ApoE ε4-dependent negative correlation between peripheral CD45RO^+^CXCR3^+^CD127^+^ CD4^+^ T cells and plasma NfL levels in patients with AD.

As a major component of adaptive immunity, CD4^+^ T cells are highly heterogeneous. Upon activation, naïve CD4^+^ T cells differentiate into effector cells with distinct cytokine and chemokine receptor profiles. For example, Th1 cells express IFN-γ and CXCR3, Th2 cells express IL-4 and CCR4, Th17 cells express IL-17 and CCR6, and Tfh cells express IL-21 and CXCR5 (*40–43*). Animal studies have demonstrated that CD4^+^ T cells are critical for maintaining cognitive function and normal behavior (*5*). Th1 cells promote cell-mediated immune responses by activating macrophages and cytotoxic CD8^+^ T cells, and are essential for host defense against viruses and intracellular bacteria and protozoa (*44*). Our transcriptomic and pathway analysis of CXCR3^+^CD127^+^ Th1 cells showed that these cells have enhanced activities of Cdc42 besides strengthened inflammation signaling. Cdc42 signaling pathway positively regulates actin cytoskeleton signaling, and facilitates immune cell activation, proliferation, migration/homing, recognition, and phagocytosis (*45*). Therefore, CXCR3^+^CD127^+^ Th1 cells appears to be highly proinflammatory, cytotoxic, proliferative, tissue homing-ready and resistant to apoptosis. Regarding the observation that plasma IFN-γ levels were not reduced in AD patients, we postulate that the lost IFN-γ production due to reduced CXCR3^+^CD127^+^ Th1 cell abundance might be compensated by other IFN-γ-producing cells with increased IFN-γ synthesis. Additionally, Th1 cells in AD patients were more activated marked by higher plasma IL-2 levels (**Supplementary Fig. 2B**), which likely rendered higher synthesis of IFN-γ per cell. Furthermore, IFN-γ can also be produced by CD8^+^ cytotoxic T cells and NK cells that were not reduced in AD patients.

In contrast to in the peripheral blood, we found that CXCR3^+^CD127^+^ Th1 cell abundance was significantly increased in the CSF of AD patients compared to healthy controls. Chemokine and chemokine receptor interaction modulates leukocyte trafficking (*46, 47*). The report of higher concentration of CXCR3 ligand CXCL10 in the CSF than that in serum (*48*) may explain our observation that a large majority of CD4^+^ T cells in CSF was CXCR3^+^CD127^+^ Th1 cells even in healthy individuals. Given aberrant elevation of CXCL10 in the CSF of AD patients (*48*), our results indicate an enhanced cell migration mediated by CXCR3-CXCL10 interaction. Moreover, our analysis showed that CXCR3^+^CD127^+^ Th1 cells in the CSF of AD patients were more activated with enhanced proinflammatory capacities compared to healthy donors. Importantly, we also found the ApoE ε4-dependent negative correlation between peripheral CXCR3^+^CD127^+^ Th1 cells and plasma NfL levels in patients with AD. ApoE ε4 is one of the most significant genetic risk factors for late-onset AD, associated with cognitive impairment and neuroinflammation (*49, 50*). Overexpression of *APOE4* decreases blood-brain barrier (BBB) integrity (*51*) and myeloid-producing apoE modulates antigen presentation to activate T cells (*52*). These observations indicate ApoE ε4 may play a role in T cell homing to the brain and activation during AD pathogenesis. NfL is a neuron specific cytoskeletal protein and its protein level increases in blood when neuronal damage occurs (*53, 54*). Although NfL is not an AD-specific biomarker, increased blood (plasma and serum) NfL levels are correlated with clinical progression in presymptomatic AD; its level is also correlated with the severity of tau pathology and neurodegeneration in post mortem brain tissue (*30–33*). Moreover, plasma NfL levels are positively associated with ApoE ε4 (*55*). Taken together, our observations suggest that homing of peripheral CXCR3^+^CD127^+^ Th1 cells to CNS potentially play a role in neuroinflammation in AD, which may also involve ApoE function.

Sex-biased differences are observed in both innate and adaptive immune responses (*56*). Emerging data have shown the impact of sex differences on AD clinical manifestation (*57, 58*). One limitation of the present study is that thirty-four out of thirty-seven samples for the CyTOF analysis were from male donors and the three remaining samples from the female donors were all in the AD group (**Supplementary Table 1**), due to the fact that ∼5% of the general veteran population are females. Additionally, there was about 10 years of age difference between two groups (**Supplementary Table 1**). To limit these two confounding factors, we applied multivariate regression models and partial correlation for statistical analysis, adjusting for age and sex, to minimize the impact of confounding on the conclusions from the CyTOF analysis on PBMC samples of the present study. We have also conducted a linear regression analysis between peripheral CXCR3^+^CD127^+^ Th1 cell frequency and various comorbidity diagnosed in these cohorts (**Supplementary Table 2**) and observed no significant correlation.

In summary, our study supports the involvement of peripheral adaptive immune cells in the pathophysiology of AD. Our findings have demonstrated that certain human CD4 T cell subpopulations of the peripheral immune system, in particular, CXCR3^+^CD127^+^ Th1 cells, are associated with AD. Further investigation is warranted to understand how these cells are involved in the modulation of neuroinflammation.

## MATERIALS AND METHODS

### Human subjects and blood samples

Participants in this study were recruited from the Dementia Care Unit at Bedford VA Hospital. The study protocol was approved by the Institutional Review Board of the VA Hospital, and written informed consent was obtained from each participant prior to the start of the study. Subjects were selected based on a clinical diagnosis of AD. All patients underwent neuroimaging, cognitive testing, and complete neurological evaluation at a memory disorder clinic. Routine blood work, including tests for thyroid-stimulating hormone (TSH) and vitamin B12, was performed on all subjects. The stage of AD dementia was classified as moderate to severe, based upon Mini-Mental State Examination (MMSE) scores ranging from 10-20 (*59*). Peripheral blood samples were obtained from patients with established AD and non-AD control at the Bedford VA Healthcare System (**Supplementary Tables 1 and 2**). Blood was collected with BD Vacutainer CPT tubes (Becton, Dickinson and Company, Franklin Lakes, NJ) and centrifuged at 1500 rcf at room temperature for 20 min. Plasma was collected and stored at –80°C. Peripheral blood mononuclear cells (PBMC) were isolated and cryopreserved in freezing media (90% fetal bovine serum, 10% dimethyl sulfoxide).

### Mass cytometry

All buffers and reagents for mass cytometry were purchased from Standard BioTools (formerly Fluidigm Corporation), unless otherwise stated. Cryopreserved PBMCs were thawed at 37 °C in a water bath and washed with RPMI (Sigma). Cells were stained with the 29-Marker Maxpar Human Immune Monitoring Panel Kit (Fluidigm, PN PRD033 C1) following the manufacturer’s instructions (see **Supplementary Table 2** for the antibody list). Briefly, cells were first stained with Cell-ID for viability followed by incubation with TruStain FcX to block non-specific Fc Receptor binding by antibodies. After washing in Maxpar Cell Staining Buffer (CSB), cells were resuspended in CSB (6×10^7^ cells/ml). An antibody cocktail (45 μl) was added to cells and the cell suspension was incubated at room temperature for 30 min. Cells then were washed with CSB twice and fixed with fixation buffer (1.6% formaldehyde in PBS) for 10 min. Cells were centrifuged and the supernatant was discarded. The cell pellet was resuspended in 1mL of cell-ID intercalator-Ir solution (125 nM) and incubated at 4 °C for overnight. On the following day, PBMCs were washed with CSB and Maxpar Cell Acquisition Solution (CAS). Cells were counted and kept at 4 °C until ready for analysis. To prepare cells for a Helios mass cytometer (Fluidigm), cells were resuspended at 1×10^6^ cells/ml in CAS containing 0.1xEQ four-element calibration beads. The Helios system equiped with CyTOF Software v6.5.358 was run using the Maxpar Direct immune profiling assay template and with WB injector to collect 50,000 300,000 events per sample, with a large majority of the samples contained > 200,000 events. Multidimensional data generated by the Helios mass cytometer was normalized with calibration beads and exported as Flow Cytometry Standard (FCS) files.

### Open-ended analysis of Mass cytometry data

The FCS files of CyTOF were analyzed with dimensionality reduction and clustering approaches using FlowJo-10 (BD Biosciences) and its plugins including DownSampleV3, Phenograph and FlowSOM. A gating strategy using the measurements of Bead, DNA1 (191Ir), Event Length, Live/dead and CD45 was applied to remove unwanted events (beads, debris, aggregates and dead cells) and select leukocytes from the FCS files of CyTOF (**Supplementary Fig. 5**). DownSampleV3 was used to randomly select equal number of events (38,000 events) from each sample and concatenated them into one file. All cell surface markers (n = 29) were selected to generate a tSNE plot (Auto, iteration 1000, perplexity 30, learning rate 28560, KNN algorism “Exact (vantage point tree)”, gradient algorithm “Barnes-Hut”). On top of the tSNE plot, the concatenated file was subjected to FlowSOM analysis with all 29 parameters for 34 cluster-clustering. Statistics on the cell proportion of each sample for each cluster were exported for calculating the statistical difference between AD patients and non-dementia (ND) controls in each cluster (two-tailed, unpaired, *t*-test) with Prism 9 (GraphPad Software), while statistics of the median staining intensity of each cell surface marker for each cluster were exported for generating heat maps for cell surface marker expression using GENE-E [http://www.broadinstitute.org/cancer/software/GENE-E/]. CD45^+^CD3^+^ live cells were selected as T cells (**Supplementary Fig. 6**) and analyzed with similar approach.

### Manual gating analysis of Mass cytometry data

After removing unwanted events in the FCS files of CyTOF using Gaussian parameters with FlowJo-10, CD45^+^ live cells were also subjected to manual gating analysis in contrast to the open-end analysis. The manual gating strategies for main cell populations for **Fig. 1C & 1D and Supplementary Fig. 1** followed manufacturer’s instruction (Fluidigm, PN PRD033 C1, Product Information Sheet) : Total T cells, CD45^+^CD3^+^; CD4 T cells, CD45^+^CD3^+^TCRγδ^−^CD4^+^CD8^−^; CD8 T cells, CD45^+^CD3^+^TCRγδ^−^CD4^−^CD8^+^; B cells, CD45^+^CD14^−^CD66b^−^CD3^−^CD19^+^; NK cells, CD45^+^CD14^−^CD66b^−^CD3^−^CD19^−^CD20^−^CD123^−^CD56^+^CD161^+^; DCs, CD45^+^CD14^−^CD66b^−^CD3^−^CD19^−^CD20^−^CD56^−^HLADR^+^; monocytes, CD45^+^CD66b^−^CD3^−^CD19^−^CD14^+^. The manual gating strategies for cell populations (clusters) that had altered cell frequencies in AD identified by the FlowSOM analysis were formed based on the antibody staining features of these clusters shown in heat maps.

### Enzyme-linked immunosorbent assay

Plasma proteins were quantified with the MesoScale Discovery Electrochemiluminescence (MSD) platform (MSD, Gaithersburg, MD) following manufacturer’s instructions. Specifically, NfL was accessed with the R-PLEX Human Neurofilament L Antibody Set. Tau and Phospho-Tau (Thr181) monoclonal antibodies BT-2, AT270 and HT-7 (Thermo Fisher Scientific, Waltham, MA) were used to capture/detect Tau and pTau181.

### Statistical analysis

Mann-Whitney test (Wilcoxon rank-sum test) was performed to compare the cell frequency (CyTOF) and protein concentration (ELISA) between AD patients and ND controls with Prism 9 (GraphPad Software). *p*-value < 0.05 was considered nominally significant. Multiple-logistic regression was conducted with *glm* function in R across the entire cohort, with AD/ND status as dependent variable, cell frequency, age and sex as independent variables, with age centered and scaled with the *scale* function. The average within group cell frequencies, 95% confidence interval and p-value of the coefficient for the cell frequency term based on standard error was reported. Partial Pearson’s correlation analysis was conducted with *pcor.test* function from the *ppcor* package in R (*60*). We reported the partial correlation coefficient within each group. Comparison and analysis of correlation coefficient between groups was carried out with the *cocor* package in R (*61*). Nominal and adjusted p-value based on Fisher’s r-to-Z transformation and Zou’s 95% confidence interval of the difference between two correlation coefficients were reported.

### Single-cell RNA-seq data analysis

Previously published single-cell RNA-seq (scRNA-seq) data of frozen PBMCs obtained from 5 healthy individuals was accessed through NCBI GEO (GSE158055) (*16*) and analyzed with Seurat v4.1.1 in R. Expression profiles of PBMCs were log-normalized to remove batch effect, scaled and clustered with 15 principal components (PCs) and a resolution of 0.5. *CD3E*^+^*CD8A*^−^*CD4*^−^*TRDC*^−^ PBMCs were identified the double negative (DN) T cell enriched cluster. Differentially expressed (DE) genes of DN T cell cluster were obtained via Wilcoxon rank-sum test comparing to other *CD3E*^+^ T cell enriched clusters and defined as having Bonferroni corrected *p*-value < 0.05. To obtain a CD4^+^ T cell-enriched population, *CD3E*^+^*CD8A*^−^*CD4*^+^*TRDC*^−^ clusters were selected and re-clustered twice with 15PCs and a resolution of 0.5, *CD8A*^+^ cluster was excluded each time to minimize CD8^+^ T cell contamination. *CXCR3*^+^*IL7R*^+^ CD4 T cell cluster was identified. DE genes of the cluster were obtained via comparison to the rest of the CD4^+^ T cells and defined as adjusted *p*-value < 0.05 from Wilcoxon rank-sum test.

scRNA-seq data of CSF isolated from 4 AD and 9 control donors were obtained from NCBI GEO (GSE134579) (*10*) with sample inclusion and exclusion criteria described in the original publication. The expression profiles were log-normalized, scaled, and effect of disease status was regressed out during the scaling process. The cells were then clustered with 15 PCs and a resolution of 0.5. *CD3E*^+^*CD8A*^−^*CD4*^+^*TRDC*^−^ clusters were selected and re-clustered with 15PCs and a 0.5 resolution. One *CD8A*^+^ cluster was excluded, the rest were re-clustered with 15PCs and a resolution of 0.5 with no significant CD8^+^ T cell contamination observed. *CXCR3*^+^*IL7R*^+^*CD4*^+^ T cells were identified and merged as one population. DE genes of the population among healthy subjects were obtained via Wilcoxon rank-sum test by comparing to the rest of the CD4^+^ T cells from healthy donors. A Bonferroni adjusted p-value < 0.05 was used to identify significant DE genes. The relative frequency of the *CXCR3*^+^*IL7R*^+^ CD4^+^ T cells among CD4^+^ T cells from AD subjects and control subjects were computed, and Chi-squared test was used to evaluate the difference of relative frequency. Phi (φ) value (sqrt(Chi-sq/n)) was used to correct for sample size, and odds ratio was used to estimate the effect size.

### Pathway Analysis

Differentially expressed genes (adjusted *p*-value < 0.05) identified in the scRNA-seq data analysis were submitted to Enrichr (http://amp.pharm.mssm.edu/Enrichr/) for transcriptional and pathway analysis (*62–64*). Input gene sets were randomly re-sampled, and Benjamini-Hochberg adjusted p-value derived from Fisher’s exact test for each replicate was used to compute a mean rank and standard deviation from the expected rank for each term in the gene-set library. The combined score is computed by taking the log of the *p*-value from the Fisher exact test and multiplying that by the z-score of the deviation from the expected rank. QIAGEN [Ingenuity Pathway Analysis](https://digitalinsights.qiagen.com/products-overview/discovery-insights-portfolio/analysis-and-visualization/qiagen-ipa/) (IPA) (Qiagen) canonical pathway analysis was performed for differentially expressed genes (adjusted *p*-value < 0.05). Canonical pathways with z-scores ≥ 2 and z-scores ≤ −2 were defined as activated and inhibited mechanisms, respectively.

## ABBREVIATIONS

AD: Alzheimer’s Disease
BCR: B-cell receptor
CAS: cell acquisition solution
CNS: central nervous system
CPT: cell preparation tube
CSB: cell staining buffer
CSF: cerebrospinal fluid
CyTOF: single-cell mass cytometry by time of flight
FCS: flow cytometry standard
IFN-γ: Interferon-γ
IL-4: Interleukin-4
IL-17: Interleukin-17
IL-21: Interleukin-21
IPA: Ingenuity Pathway Analysis
KNN algorism: k-nearest neighbors algorithm
MCI: mild cognitive impairment
MS: multiple sclerosis
ND: non-dementia
PBMC: peripheral blood mononuclear cell
PD: Parkinson’s disease
RhoGDI: Rho GDP dissociation inhibitor
scRNA-seq: single-cell RNA-sequencing
SOM: self-organizing map
TCR: T-cell receptor
TRAC: T cell receptor alpha constant
TRDC: T cell receptor delta constant
tSNE: t-distributed stochastic neighbor embedding

## DECLARATIONS

### Ethical approval and consent to participate

The study was approved by the Bedford VA Healthcare System Institutional Review Board, and the written informed consent was obtained from all participants. All procedures were performed according to the National Institutes of Health’s guidelines for research using human subjects.

## ACKNOWLEDGEMENTS

We thank Dr. Lauren Moo for helpful discussion. The views expressed in this article are those of the authors and do not represent the views of the US Department of Veterans Affairs or the US Government.

## Author contributions

D.H. designed and supervised the study, analyzed data and wrote the manuscript; M.C. designed and performed experiments; X.L. conducted data analysis and wrote the manuscript; S.D. performed experiments; P.M. recruited study subjects and wrote the manuscript; Y.H. and M.H. conducted data analysis and edited the manuscript. H.L.W. supervised the study, analyzed data and wrote the manuscript; W.X. designed and supervised the study, analyzed data and wrote the manuscript.

## Data and materials availability

The authors declare that the main data supporting the findings of this study are available within the article and its [Supplementary Information](https://www.nature.com/articles/ncomms12998#s1) files. Additional data are available from the corresponding authors upon request.

