## Supplementary tables and figures for "ApoE ε4-dependent alteration of CXCR3^+^CD127^+^ CD4^+^ T cells is associated with elevated plasma neurofilament light chain in Alzheimer’s disease"

**List of Supplementary Materials**

**Supplementary Tables:**

Supplementary Table 1 Demographics for AD patients and non-AD controls.

Supplementary Table 2 ApoE type and cohort comorbidities.

Supplementary Table 3 Antibody panel for immune cell surface markers.

Supplementary Table 4 No significant changes in the cell frequency of double negative T cells and $CXCR3^{+}CD127^{+}$ Th1-subset among patients with AD.

Supplementary Table 5 Partial Pearson's correlation coefficient of cell population frequency among T cells versus NfL levels controlling for age and sex.

Supplementary Table 6 Partial Pearson's correlation coefficient of cell population frequency in PBMCs versus NfL levels controlling for age, sex and *APOE* type.

**Supplementary Figures:**

Supplementary Figure 1 Manual gating analysis reveals no significant difference in the primary immune cell types between AD patients and non-dementia donors.

Supplementary Figure 2 Plasma IFN-γ and IL-2 levels in AD patients.

Supplementary Figure 3 Scheme for extracting the gene expression profiles of peripheral $CD4^+$ T cell-enriched populations.

Supplementary Figure 4 Scheme for extracting the gene expression profiles of CSF $CD4^+$ T cell-enriched populations.

Supplementary Figure 5 Preprocessing gating strategy for CyTOF dataset

Supplementary Figure 6 Gating strategy for total T cells.

### SUPPLEMENTARY MATERIALS

### SUPPLEMENTARY TABLES

**Supplementary Table 1** Demographics for AD patients and non-AD controls.

| Group (# of subjects) | | Total (N=37) | AD (N=18) | Non-AD (N=19) |
|---|---|---|---|---|
| | | Mean (SD) | Mean (SD) | Mean (SD) |
| **Age (years)** | | 73.6 (9.1) | 78.1 (10.5) | 69.3 (4.9) |
| | | N (%) | N (%) | N (%) |
| **Sex (#)** | Female | 3 (8.1%) | 3 (16.7%) | 0 (0%) |
| | Male | 34 (91.9%) | 15 (83.3%) | 19 (100%) |
| **Race (#)** | White | 25 (67.6%) | 12 (66.7%) | 13 (68.4%) |
| | American Indian or Alaska Native | 0 (0%) | 0 (0%) | 0 (0%) |
| | Asian | 0 (0%) | 0 (0%) | 0 (0%) |
| | Black or African American | 0 (0%) | 0 (0%) | 0 (0%) |
| | Native Hawaiian or Other Pacific Islander | 0 (0%) | 0 (0%) | 0 (0%) |
| | Unassigned | 12 (32.4%) | 6 (33.3%) | 6 (31.6%) |

**Supplementary Table 2** ApoE type and cohort comorbidities.

Note: na, not available. 1, positive diagnosis.

| Sample ID | Group | Age | Sex | Sex | Comorbidities | | | | | | |
|---|---|---|---|---|---|---|---|---|---|---|---|
| | | | | ApoE | Diabetes | Hypertension | Heart Failure | Kidney Disease | Hyperlipidemia | Cancer | Stroke |
| 7 | Control | 68 | Male | ε2/ε3 | | | | | | | |
| 8 | Control | 63 | Male | ε3/ε3 | 1 | 1 | 1 | 1 | | 1 | 1 |
| 9 | Control | 69 | Male | ε3/ε3 | | | 1 | | 1 | | |
| 10 | Control | 65 | Male | ε3/ε4 | 1 | | | | 1 | | |
| 12 | Control | 66 | Male | ε3/ε4 | 1 | | | | | | |
| 13 | Control | 64 | Male | ε3/ε3 | | | | | 1 | | |
| 19 | Control | 70 | Male | ε3/ε3 | 1 | 1 | | | 1 | | |
| 20 | Control | 68 | Male | ε2/ε2 | 1 | | | | | | |
| 26 | Control | 69 | Male | na | | | | 1 | 1 | | |
| 29 | Control | 77 | Male | ε3/ε3 | 1 | | | 1 | | | |
| 30 | Control | 75 | Male | ε3/ε3 | | | | | | | |
| 31 | Control | 68 | Male | ε3/ε3 | | | | | 1 | | 1 |
| 35 | Control | 70 | Male | ε3/ε3 | | | | | | | |
| 41 | Control | 67 | Male | ε3/ε4 | | | | | | | |
| 42 | Control | 68 | Male | ε2/ε4 | | | | | | | |
| 49 | Control | 70 | Male | ε3/ε3 | | | | | | | |
| 51 | Control | 84 | Male | ε2/ε3 | | | | | | | |
| 59 | Control | 66 | Male | ε3/ε3 | | | | | | | |
| 69 | Control | 70 | Male | ε3/ε3 | 1 | 1 | | | 1 | | 1 |
| 2 | AD | 66 | Male | na | | | | | | | |
| 3 | AD | 61 | Female | ε3/ε3 | | | | | | | |
| 4 | AD | 90 | Female | ε3/ε3 | | 1 | | | | | |
| 5 | AD | 62 | Male | ana | | 1 | | | | | |
| 22 | AD | 67 | Male | ε3/ε4 | | 1 | | | | | |
| 33 | AD | 83 | Male | ε3/ε4 | | | | | | | |
| 38 | AD | 87 | Male | ε3/ε4 | | 1 | | | | | |
| 39 | AD | 80 | Male | ε3/ε4 | | | | | | | |
| 40 | AD | 66 | Male | ε3/ε4 | 1 | | | | 1 | | |
| 43 | AD | 85 | Male | ε3/ε4 | | 1 | | | | | |
| 44 | AD | 70 | Male | ε3/ε4 | | | | | | | |
| 48 | AD | 89 | Male | ε3/ε4 | | 1 | | | | | |
| 53 | AD | 73 | Male | ε3/ε4 | | | | | | | |
| 54 | AD | 92 | Female | ε3/ε4 | | | | | | | |
| 56 | AD | 84 | Male | ε3/ε3 | | | | | | | |
| 63 | AD | 90 | Male | ε3/ε3 | | | | | | | |
| 64 | AD | 85 | Male | ε3/ε4 | | | | | | | |
| 65 | AD | 73 | Male | ε4/ε4 | | | | | | | |

**Supplementary Table 3** Antibody panel for immune cell surface markers.

| Antigen | Clone | Metal |
|---|---|---|
| CD3 | UCHT1 | 154Sm |
| CD4 | SK3 | 174Yb |
| CD8 | SK1 | 168Er |
| CD11c | Bu15 | 147Sm |
| CD14 | M5E2 | 175Lu |
| CD16 | 3G8 | 148Nd |
| CD19 | HIB19 | 142Nd |
| CD20 | 2H7 | 171Yb |
| CD24 | ML5 | 166Er |
| CD25 | 2A3 | 169Tm |
| CD27 | L128 | 158Gd |
| CD28 | CD28.2 | 160Gd |
| CD38 | HIT2 | 144Nd |
| CD45 | HI30 | 89Y |
| CD45RA | HI100 | 155Gd |
| CD45RO | UCHL1 | 165Ho |
| CD56 | NCAM16.2 | 176Yb |
| CD66b | 80H3 | 162Dy |
| CD123/IL-3R | 6H6 | 151Eu |
| CD127/IL-7Rα | A019D5 | 143Nd |
| CD161 | HP-3G10 | 164Dy |
| CD183/CXCR3 | G025H7 | 163Dy |
| CD185/CXCR5 | RF8B2 | 153Eu |
| CD194/CCR4 | L291H4 | 149Sm |
| CD196/CCR6 | G034E3 | 141Pr |
| CD197/CCR7 | G043H7 | 167Er |
| HLA-DR | L243 | 173Yb |
| IgD | IA6-2 | 146Nd |
| TCRγδ | 11F2 | 152Sm |

**Supplementary Table 4** No significant changes in the cell frequency of double negative T cells and $CXCR3^{+}CD127^{+}$ Th1-subset among patients with AD.

Multivariate regression model of the cell frequencies (%) of double negative T cells and $CXCR3^{+}CD127^{+}$ Th1-subset within PBMCs, adjusted for age and sex.

| Cell subset | AD (%) | ND (%) | 95% CI | *p*-Value |
|---|---|---|---|---|
| $CD4^{-}CD8^{-}$ (DN) | 0.39 | 0.63 | -0.370 to 0.091 | 0.225 |
| $CXCR3^{+}CD127^{+}$ (Th1-subset) (of PBMCs) | 5.46 | 8.75 | -6.424 to 0.697 | 0.111 |

**Supplementary Table 5** Partial Pearson's correlation coefficient of cell population frequency among T cells versus NfL levels controlling for age and sex.

| | Correlation | | AD | | |
|---|---|---|---|---|---|
| Cell subset | AD | ND | 95% CI | *p*-Value | Adj *p*-Value |
| Th2-subset | 0.25 | -0.22 | (-0.21, 1.03) | 0.18 | 0.4 |
| cmCD4-subset | 0.15 | 0.058 | (-0.57, 0.72) | 0.81 | 0.81 |
| Th17-subset | -0.43 | -0.0006 | (-0.99, 0.22) | 0.2 | 0.4 |
| Th1-subset | -0.19 | 0.065 | (-0.86, 0.43) | 0.48 | 0.64 |

**Supplementary Table 6** Partial Pearson's correlation coefficient of cell population frequency in PBMCs versus NfL levels controlling for age, sex and ApoE type.

| Cell subset | Correlation AD | Correlation ND | AD 95% CI | AD *p*-Value |
|---|---|---|---|---|
| Th2-subset (of PBMCs) | 0.091 | -0.043 | (-0.57, 0.79) | 0.72 |
| Th1-subset (of PBMCs) | -0.71 | 0.078 | (-1.27, -0.17) | 0.011 |

### SUPPLEMENTARY FIGURES AND FIGURE LEGEND

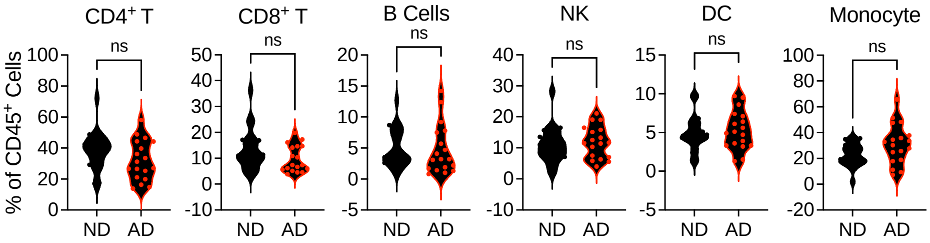


**Supplementary Figure 1 Manual gating analysis reveals no significant difference in the primary immune cell types between AD patients and non-dementia donors.**

The cell frequency of $CD4^+$ T cells, $CD8^+$ T cells, B cells, NK cells, dendritic cells, monocytes was compared between non-dementia controls (ND, n = 19) and AD patients (AD, n = 18). ns, not significant (Mann-Whitney test, mean ± s.d.).

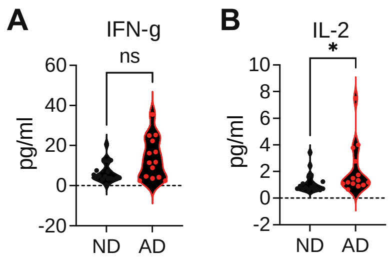


**Supplementary Figure 2 Plasma IFN-γ and IL-2 levels in AD patients.**

The plasma levels of IFN-γ and IL-2 were quantified by ELISA. ns, not significant; *, *p* value < 0.05 (Mann-Whitney test, mean ± s.d.).

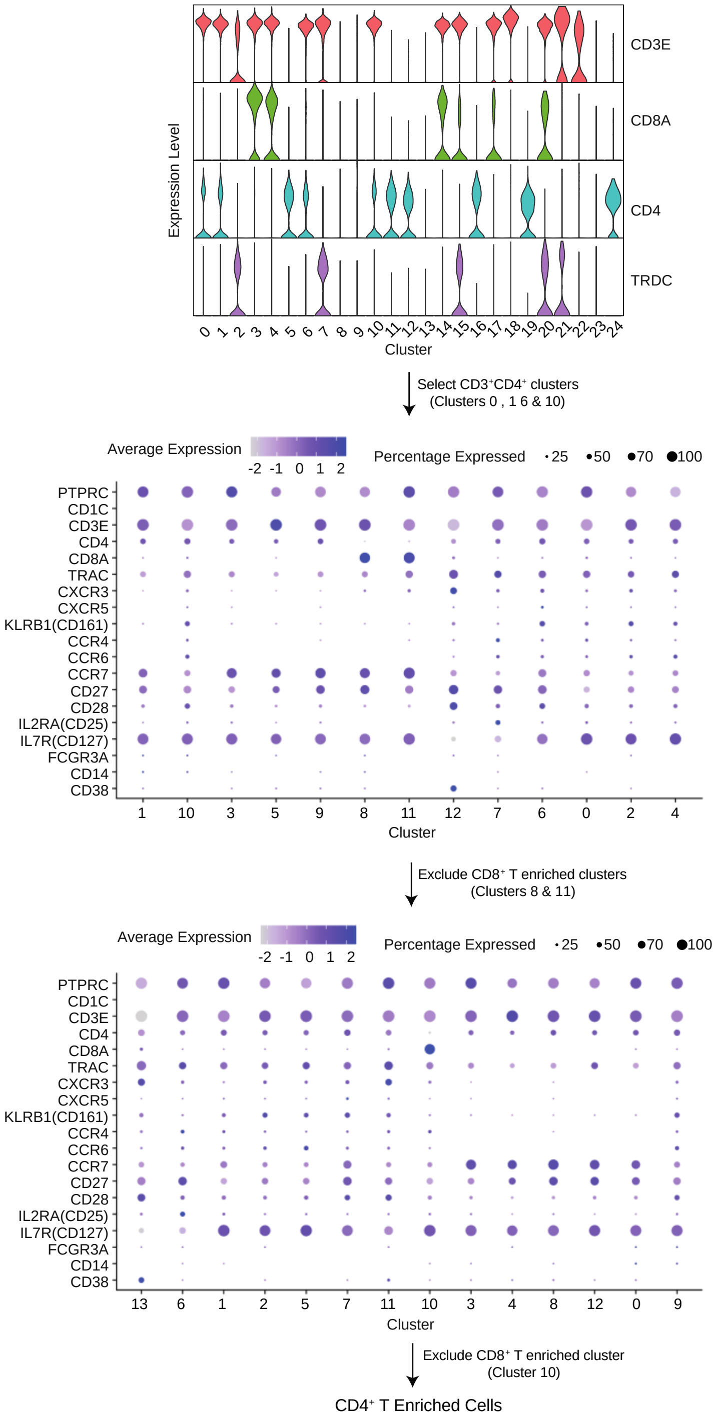


**Supplementary Figure 3 Scheme for extracting the gene expression profiles of peripheral CD4$^+$ T cell-enriched populations.**

Expression profiles of $CD4^+$ T cells were extracted from a public single-cell RNA-seq (scRNA-seq) dataset (GSE158055). Briefly, PBMCs were clustered into 25 clusters and *CD3E*, *CD8A*, *CD4* and *TRDC* expression in clusters was shown (upper panel). Cells in *CD3E*$^+$*CD8A*$^-$*CD4*$^+$*TRDC*$^-$ clusters were selected and re-clustered to into 13 clusters (middle panel). Two *CD8A*$^+$ clusters were selected out. The remaining cells were further clustered into 14 clusters (lower panel). One *CD8A*$^+$ cluster was selected out. The remaining cells formed the CD4 T cell-enriched population for further analysis.

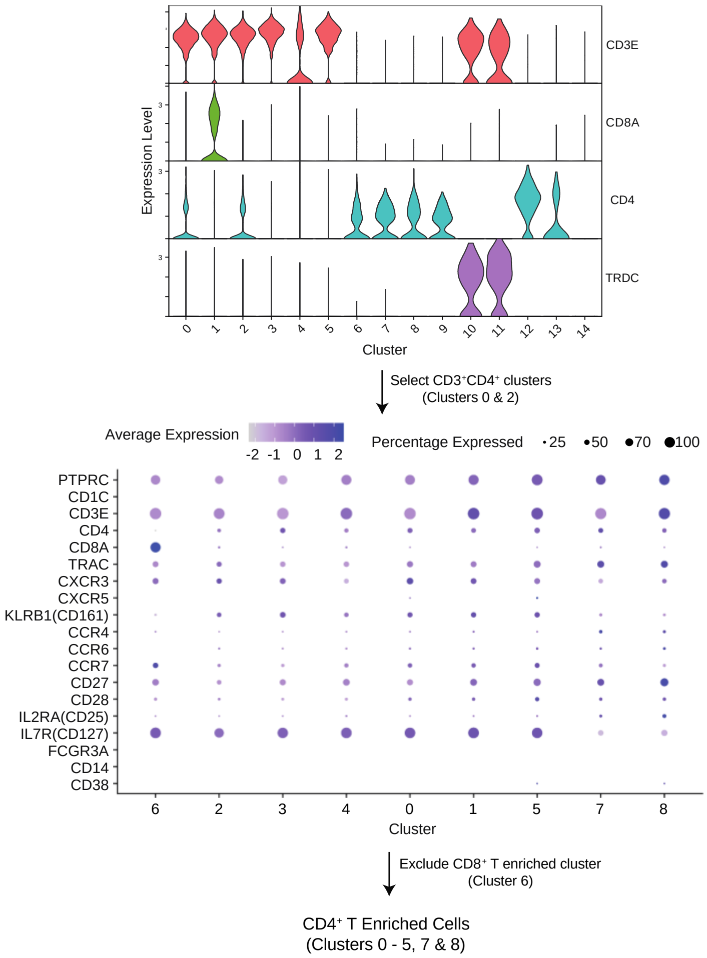


**Supplementary Figure 4 Scheme for extracting the gene expression profiles of CSF CD4$^+$ T cell-enriched populations.**

Expression profiles of CSF CD4$^+$ T cells were extracted from a public single-cell RNA-seq (scRNA-seq) dataset on CSF cells from AD patients and healthy controls (GSE134579). Briefly, CSF cells were clustered into 15 clusters and *CD3E*, *CD8A*, *CD4* and *TRDC* expression in clusters was shown (upper panel). Cells in *CD3E*$^+$*CD8A*$^-$*CD4*$^+$*TRDC*$^-$ clusters were selected and re-clustered to into 9 clusters (lower panel). One *CD8A*$^+$ cluster was selected out. The remaining cells formed the CSF CD4$^+$ T cell-enriched population for further analysis.

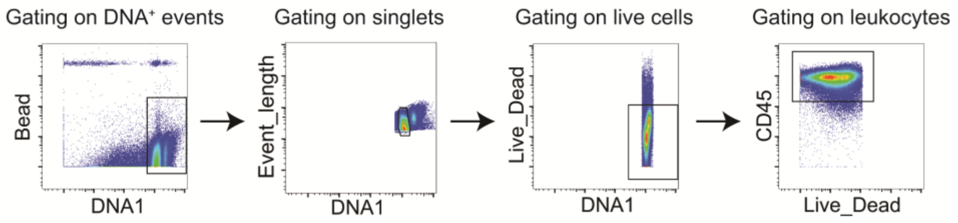


**Supplementary Figure 5 Preprocessing gating strategy for CyTOF dataset**

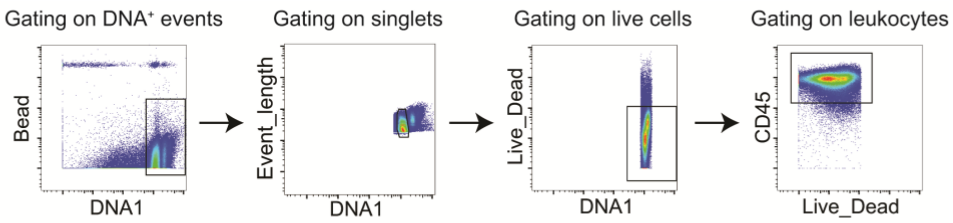


**Supplementary Figure 6 Gating strategy for total T cells.**
